# Modelling and measuring effects of shear stress in extrusion bioprinting of endothelial- epithelial cell co-cultures

**DOI:** 10.64898/2026.09.02.748801

**Authors:** Jordan W. Davern, Angus Weekes, Jorge Alberto Amaya Catano, Christoph Meinert, Laura J. Bray, Travis J. Klein

## Abstract

Extrusion-based bioprinting enables the development of tissue-like constructs; however, the impact of printing-associated shear stress on cell viability and function remains a critical consideration. To address this, we developed a comprehensive workflow combining rheological characterization, computational fluid dynamics (CFD) modelling, and experimental validation to predict and assess shear stress effects during bioprinting. The rheological properties of gelatin methacryloyl (GelMA) at 5 % (w/v, 20 °C) and 10 % (30 °C) concentrations were modelled, comparing various non-Newtonian regression models. CFD simulations were validated using micro-particle image velocimetry, showing agreement between predicted and measured velocities. The impact of bioprinting-associated shear stress on cell viability was assessed using a co-culture of human umbilical vein endothelial cells and breast epithelial cells. Immediate post-printing analysis revealed increased apoptosis in GelMA 5 % (w/v, 20 °C), although 10 % (w/v) GelMA demonstrated higher shear stress levels compared to 5 % GelMA. After 1 day of culture in crosslinked hydrogels, apoptosis increased in extrusion pressure, demonstrating the impact of low levels of acute shear stress. This workflow provides a robust methodology for predicting acute shear stress impacts during bioprinting, laying the foundation for future optimization studies.

## 1.0 Introduction

Three-dimensional (3D) bioprinting is a form of biofabrication that enables the precise fabrication of biologically organized constructs using cells, biomaterials, and bioactive molecules [1–3]. Extrusion-based bioprinting is one of the most popular biofabrication techniques that involves the layer-by-layer deposition of biomaterial-cell suspensions known as bioinks through a nozzle using pneumatic, piston-driven, or screw-driven mechanisms [2,4,5]. To develop physiologically relevant tissue-like constructs with precise patterning, materials must exhibit steady flow during printing and stabilize once deposited [2,3,6]. Biomaterials routinely used in 3D cell culture have been translated into bioink formulations such as alginate, hyaluronic acid, gelatin methacryloyl (GelMA) and polyethylene glycol (PEG), relying on controlling a material’s thermal properties or crosslinking (ionic or visible light) for rapid stabilization [5–8]. Many traditional hydrogel-based materials have been modified to possess specific viscoelastic characteristics to facilitate smooth extrusion, maintain structural fidelity during printing, and provide a suitable microenvironment for cellular growth and proliferation [1,2,4]. Rheological analysis is a critical tool utilized during bioink development, focusing on key rheological parameters, and providing insights into the prospective material’s printability [9–11]. This analysis can include evaluating viscosity and its shear-thinning capacity, viscoelasticity during filament deposition, thermal sensitivity, and gel formation (crosslinking kinetics) [8,10,12–15].

However, bioinks can induce shear stress on encapsulated cells during the extrusion process due to material properties, geometric factors, and printing conditions [16–20]. The extrusion pressure required for bioink deposition is directly proportional to the material’s viscosity; higher viscosity bioinks require greater extrusion pressures, subjecting the encapsulated cells to elevated shear stresses affecting cell viability, proliferation, and functionality in printed constructs [5,21,22]. Cells *in vivo* are exposed to a gradient of shear stress levels (∼ 0.01 – 9.5 Pa) dependent upon their location [16,17,23,24]. While excessive shear stress is considered detrimental, shear stress within the physiological range promotes cellular maturation, differentiation, and structural alignment [16]. Several studies have investigated the effects of shear stress on various primary human cell types, cell lines and animal-derived cells including human mesenchymal stromal cells (hMSCs), skin-derived fibroblasts, mammary epithelial cells, human umbilical vein endothelial cells (HUVECs), breast cancer cell lines (MDA-MB 231 and MCF-7) and mouse embryonic stem cells (ESCs) [19,21–23,25–28]. Depending on the material, nozzle geometry and cell type, various shear stress levels have been reported to impact their viability and functionality. For instance, hMSCs encapsulated in 0.5-1.5 % (w/v) alginate were affected by higher shear stress (5,000-10,000 Pa) during micro-valve extrusion, human dermal fibroblasts in a composite material consisting of 2 % (w/v) fibrinogen, 5 % (w/v) alginate and 5 % gelatin demonstrated 90-94 % viability of cells when exposed to maximum wall shear stress of 3,890-4,619 Pa within a 400 µm conical nozzle [21,25,29]. However, reduced nozzle geometries (200-100 µm) increased maximum wall shear stress to 4,789-7,002 Pa, resulting in a decrease of cell viability (73-39 %) [29].

Several studies have selected alginate alone or mixed with different polymers such as gelatin for shear stress studies due to its wide use within the tissue engineering community [20,21,27,28,30]. While alginate is shear-thinning and highly customizable, it lacks cell-binding motifs such as Arginylglycylaspartic acid (RGD) sequences, limiting cell adhesion sites and tissue-specific functions compared to collagen or gelatin-based materials that can mimic the native extracellular matrix (ECM) and tissue-specific microenvironment [2,5,31]. This distinction becomes increasingly important as bioprinting moves from single-cell-type printed constructs toward co-culture systems that recapitulate stromal, epithelial, endothelial, or disease- microenvironment interactions. Recent bioprinted co-culture models have demonstrated the value of spatially organizing chondrocyte/stem-cell combinations, endothelial-cell-containing pre-vascular structures, and multicellular cancer models, including systems incorporating HUVECs with stromal or tumor-associated cells [32–35] .Despite this progress, current shear-stress studies remain weighted toward single-cell cultures, leaving a gap in understanding how extrusion-associated shear affects co-cultures composed of multiple cell types.[9,16,36–38].

Computational fluid dynamics (CFD) has become an effective technique to simulate and predict bioprinting processes such as shear stress, velocity profiles and flow dynamics [20,39–42]. The incorporation of CFD simulations has enabled researchers to refine printing parameters and nozzle geometries, accelerating experimentation and development of bioprinting processes [43–46]. However, many studies lack proper validation against experimental data or application of non-Newtonian fluid models that accurately model the complex rheological behavior of bioinks [26,47–49]. Several studies have investigated the impact of bioprinting on *in vitro* models utilizing CFD simulations, examining various aspects such as hydrostatic pressure on HUVECs in alginate [28] shear stress on skin-derived fibroblasts across different nozzle geometries [29], and the mixing of bioink components [49]. However, very few studies have evaluated various non-Newtonian models for shear stress based on rheological data. Furthermore, limited studies have focused on validating CFD simulations against experimental flow data, which is critical for accurately representing simulated bioprinting processes.

In this study, we hypothesized that shear stress would influence apoptosis and cell viability in a HUVEC-MCF-10A co-culture. Here, we demonstrate a validated CFD workflow that incorporates rheological data for modeling shear stress in prospective bioinks and co-culture proof of concept. GelMA was selected as the model bioink due to its versatility in 3D cell culture and its ability to support various cell processes. GelMA’s flow behavior using rheometry and non-Newtonian regression models was characterized. Experimental printing parameters were collected between 20-200 kPa extrusion pressure and input into ANSYS® Fluent CFD software, with simulations validated through Micro-particle image velocimetry (micro-PIV) measurements. These models were used to investigate the immediate and acute effect of bioprinting on HUVEC/MCF-10A co-culture using 50-200 kPa extrusion pressure. The approach developed in this study provides a proof-of-concept for predicting shear stress-induced effects on cell viability and function in extrusion-based bioprinting applications.

## Results

To simulate bioprinting processes and visualize shear stress, we first conducted rheological characterization of GelMA. This involved analyzing their viscosity profiles to assess shear-thinning properties and compare non-Newtonian regression models, which will be used as inputs in computational fluid dynamics (CFD) simulations.

GelMA hydrogel systems may be modified by altering polymer concentration for modelling tissue-specific microenvironments based on tunable stiffness. However, different polymer concentrations exhibit distinct flow properties and stability at different temperatures (Figure 1A). GelMA concentrations of 5 % (w/v) at 20 °C and 10 % at 30 °C were selected to demonstrate their effect on flow behavior and simulated shear stress. By controlling the temperature of 5-10 % (w/v) GelMA, an increase in shear-thinning behavior was observed compared to their Newtonian-like behavior at 37 °C. Cooling 5 % (w/v) GelMA from 37 °C to 20 °C increased the initial viscosity from an average of 0.0201 to 0.078 Pa·s. By contrast, 10 % (w/v) GelMA showed minimal viscosity change between 37 °C and 30 °C, with average viscosities ranging from 0.0147 to 0.0217 Pa·s (Figure 1A, i). These conditions therefore provided two GelMA systems with different flow behavior for downstream shear-stress modeling.

**Figure 1:**
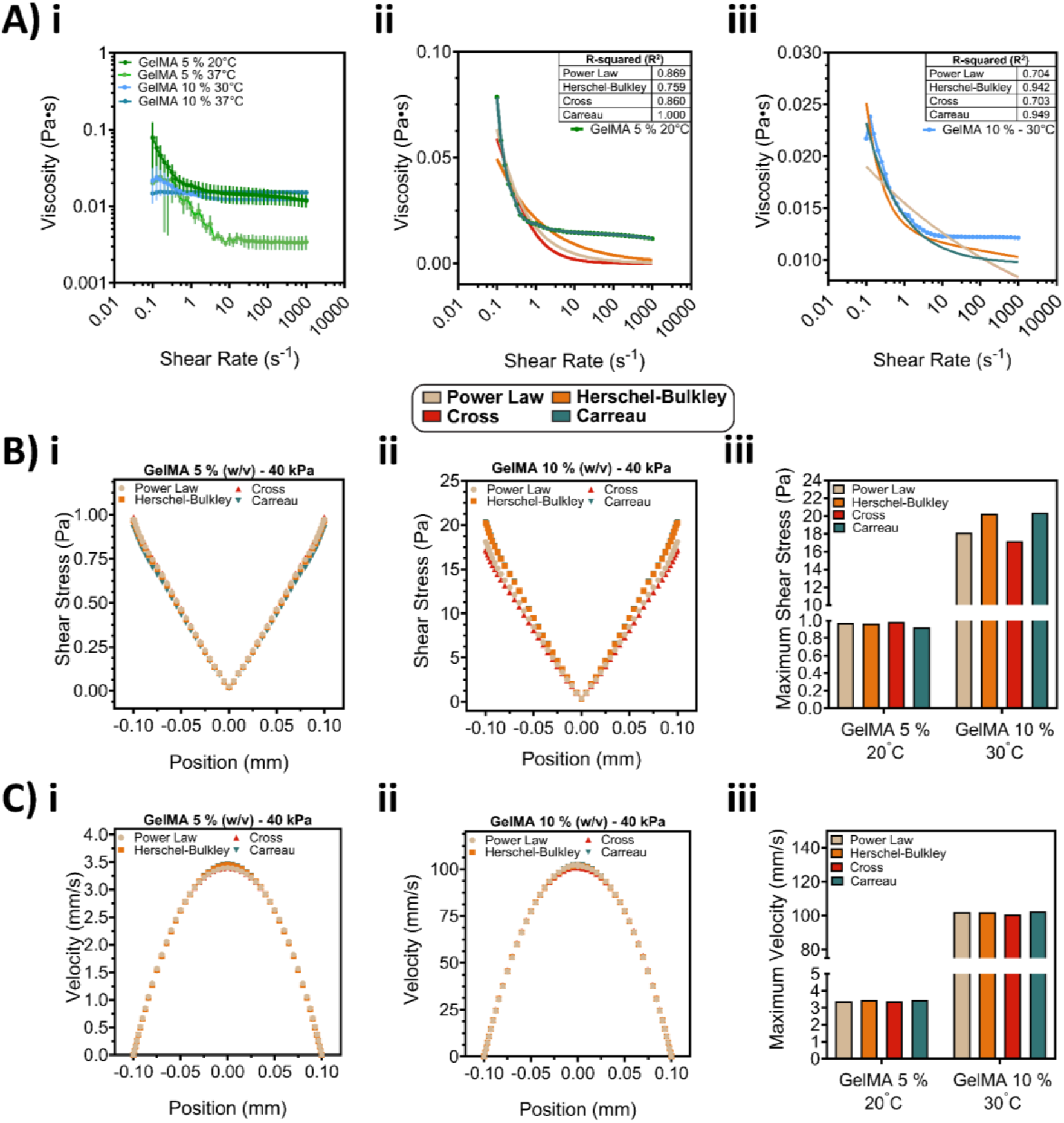
Comparison of non-Newtonian regression models using GelMA for computational fluid dynamic modelling. A) i) Temperature-controlled viscosity shear rate profile of 5 % (20 °C and 37 °C) and 10 % (w/v) (30 °C and 37 °C) GelMA, ii-iii) non-Newtonian regression model comparison of 5 % and 10 % (w/v) GelMA. B) i-ii) Simulated shear stress (Pa) profile at 27G nozzle outlet (200 µm) of 5 % and 10 % (w/v) GelMA using defined extrusion pressure (40 kPa), iii) Maximum shear stress (Pa) of outlet profile. B) i-ii) Simulated velocity (mm/s) profile at 27G nozzle outlet (200 µm) of 5 % and 10 % (w/v) GelMA using defined extrusion pressure (40 kPa), iii) Maximum velocity (mm/s) of outlet profile. The mean values and error bands representing standard deviation (SD) are shown and sample size (n) = 5 for A) i). Statistical analysis was performed for A) ii-iii) using linear regression for best fit and R2. All simulations utilize an axisymmetric model, outlet profiles of shear stress (B) i-ii) and velocity (C) i-ii) are mirrored to represent the developed outlet profiles.

Non-Newtonian regression models were compared for GelMA 5 % and 10 % (w/v) to assess best fit and R^2^ values (Equation 1-4). The Power Law, Herschel-Bulkley and Cross models demonstrated similar flow behavior with *n* indices (Newtonian fluids =1, shear thinning < 1, and shear-thickening >1) of 0.322-0.327 for GelMA 5 % (w/v) at 20 °C and 0.857-0.974 for GelMA 10 % w/v at 30 °C (Supplementary Data, Table 1). The Carreau model provided the strongest overall fit, with R^2^ values of0.949-1.0 across both concentrations, and better captured viscosity across measured shear-rate range. The other regression models fit the experimental data within the linear region at lower shear rates (0.1-1 s^-1^), but underestimated viscosity at higher shear rates (1-1000 s^-1^) (Figure 1A, ii-iii).

The non-Newtonian models were further evaluated using ANSYS® Fluent at a fixed extrusion pressure of 40 kPa for each GelMA concentration to assess the influence of regression models on simulated shear stress and velocity profiles. The simulated shear stress profiles at the 27G conical nozzle outlet showed minimal differences between regression models for GelMA 5 % (w/v) at 20 °C, with ∼3 Pa difference observed between models for the same concentration and temperature (Figure 1B, i-iii). Similarly, the velocity profiles and maximum velocities for both GelMA concentrations exhibited minimal variation (Figure 1C, i-iii). Comparison of regression models revealed that the Carreau model demonstrated superior fit and higher R² values, effectively capturing GelMA’s rheological behavior, it was selected for subsequent simulations.

Mass flow rate was next measured across 20-200 kPa extrusion pressure and modeled using MODDE® design of experiments to systematically investigate the relationship between extrusion pressure and mass flow rate for all materials. Water and GelMA 10 % (w/v) at 30 °C demonstrated near-linear increase in mass flow with increased pressure, whereas GelMA 5 % (w/v) at 20 °C produced lower mass flow rates, consistent with its temperature-dependent viscosity (Figure 2A, i).

**Figure 2:**
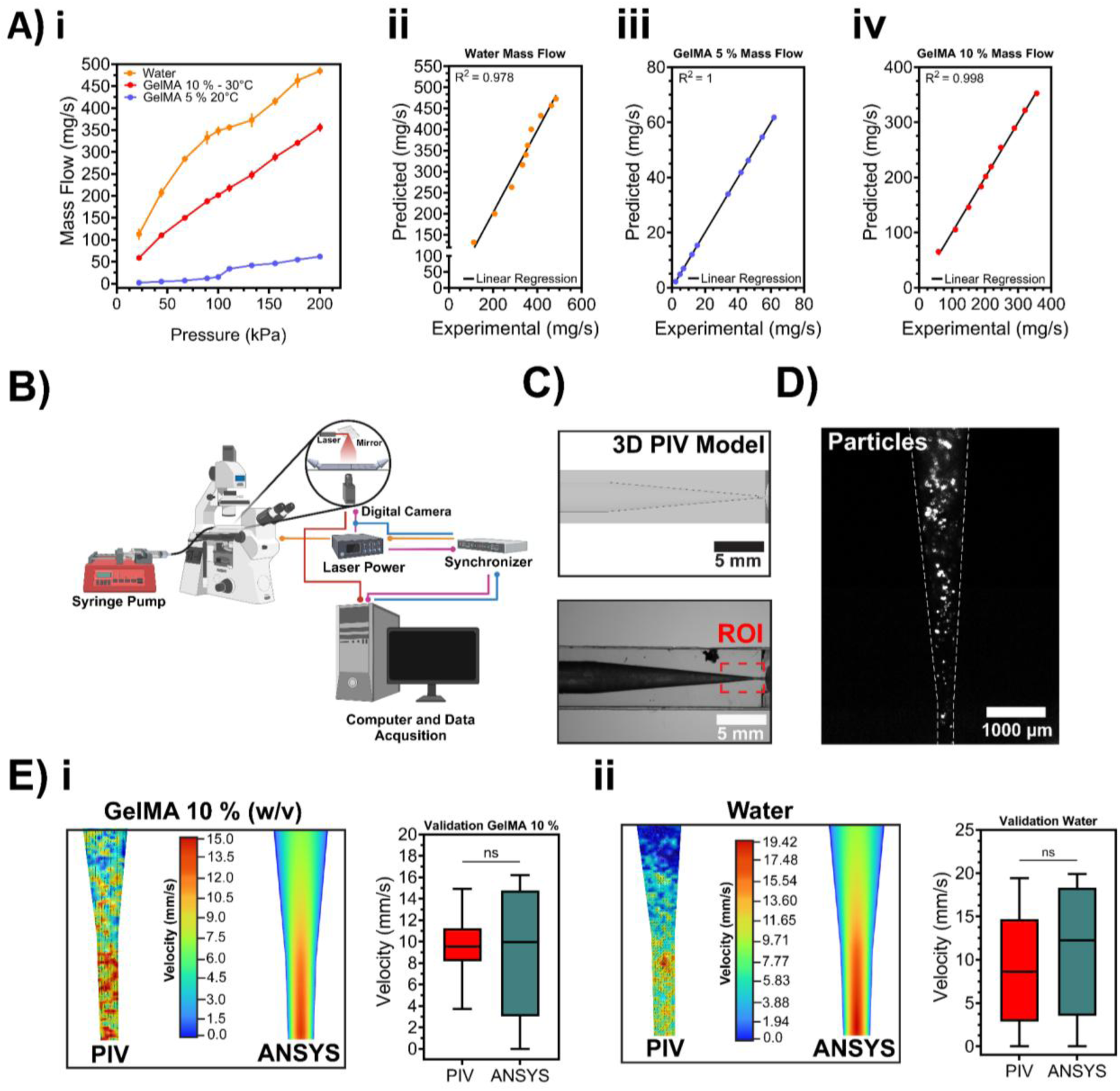
Experimental validation of computational fluid dynamics models. A) i) Mass flow rate correlation at printable extrusion pressures (20-200 kPa) of water, GelMA 5 % (w/v) – 20 °C and GelMA 10 % (w/v) – 30 °C (n = 5), ii-iv) Linear regression analysis comparing predicted mass flow rate (mg/s) vs. experimental mass flow rate of water, GelMA 5 % (w/v) – 20 °C and GelMA 10 % (w/v) – 30 °C at an extrusion pressure range of 20-200 kPa. B) Micro-PIV laser and optic configuration, created in BioRender. C) 3D CAD model of 27G nozzle for micro-PIV and printed model, red dashed lines represent micro-PIV region of interest. D) Representative frame of fluorescent particles during micro- PIV acquisition. E) i-ii) Validation of CFD simulation comparing micro-PIV velocity (mm/s) of GelMA 10 % (w/v) – 30 °C material and water. Mean values with error bands representing standard deviation (SD), statistical analysis was performed for A) i-iv) using linear regression for best fit and R^2^, E) i-ii) unpaired t-test with Welch’s correction, ns = non-significant. All simulations utilize an axisymmetric model, contour profiles of shear stress (E) i-ii) are mirrored for accurate representation.

The predicted flow rates obtained from MODDE® demonstrated a strong linear relationship with the experimental data for water and both GelMA formulations (5 and 10 % w/v) with R^2^ values of 0.978-1.0 and normally distributed residuals (Figure 2A i-iii, Supplementary Figure 3A i-iii). Narrow 95 % confidence interval bands, supported a predominately linear relationship between increasing mass flow (mg/s) with extrusion pressure (kPa) (Supplementary Data Figure 3B, i-iii). Subsequently, the models were employed in all upcoming CFD simulations, where predicted mass flow was used to calculate velocity for the inlet conditions.

To validate the velocity observed in CFD simulations, micro-PIV analysis was conducted using a 3D printed phantom model of 27G nozzle (Supplementary Data Figure 6). Micro-PIV analysis measured the velocity of fluorescent particles in two different materials: water and GelMA 10 % (w/v) at 30 °C (Figure 2B-D). Micro-PIV measured mean velocities of 9.7 ± 2.3 mm/s for GelMA 10 % (w/v) and 8.8 ± 6.1 mm/s for water, while CFD predicted 8.9 ± 5.8 mm/s and 11.1 ± 7.3 mm/s, respectively (Figure 2E i-ii). Velocities measured by micro-PIV and predicted by CFD were not significantly different (Figure 2E i-ii), supporting use of the model for downstream shear-stress interpretation.

The influence of bioprinting-associated shear stress on cell viability was investigated using HUVECs and MCF-10A cells. Prior to crosslinking cell responses were assessed in GelMA 5 % (w/v) at 20 °C and 10 % at 30 °C at 50-200 kPa. CFD simulations predicted maximum outlet shear stresses of 10–18 Pa across the GelMA 5% (w/v) printing conditions, whereas substantially higher outlet shear stresses of 55–73 Pa were predicted for GelMA 10 % (w/v) (Figure 3A-C). In GelMA 5 % (w/v), live/healthy (Caspase 3/7⁻/PI⁻) cells predominated, ranging from 86.8 % to 98.6 % across the manual pipetting control (20 °C) and extrusion pressures of 50–200 kPa (Figure 3D). Increasing extrusion pressure was associated with reduced apoptotic (Caspase 3/7⁺/PI⁺) and dead (PI⁺) cell populations. Consistent with these observations, median Caspase 3/7 fluorescence was greatest in the 50 kPa group, while the remaining groups exhibited comparable fluorescence intensities (Figure 3F). In GelMA 10 % (w/v) (30 °C), cell populations were broadly comparable between manual and printed groups. Across extrusion pressures of 50-200 kPa, apoptotic and dead cell populations remained low, ranging from 0.6-0.9 % and 1.0-1.6 %, respectively (Figure 3 E-G). Caspase 3/7 fluorescence was similarly elevated in the 50 kPa group, whereas PI fluorescence increased progressively from 50 to 200 kPa (Figure 3G). Despite the substantially higher simulated outlet shear stresses in GelMA 10 %, the greatest immediate apoptotic response occurred in the GelMA 5 % 50 kPa group, indicating that acute cell responses were not explained by peak shear stress alone.

**Figure 3:**
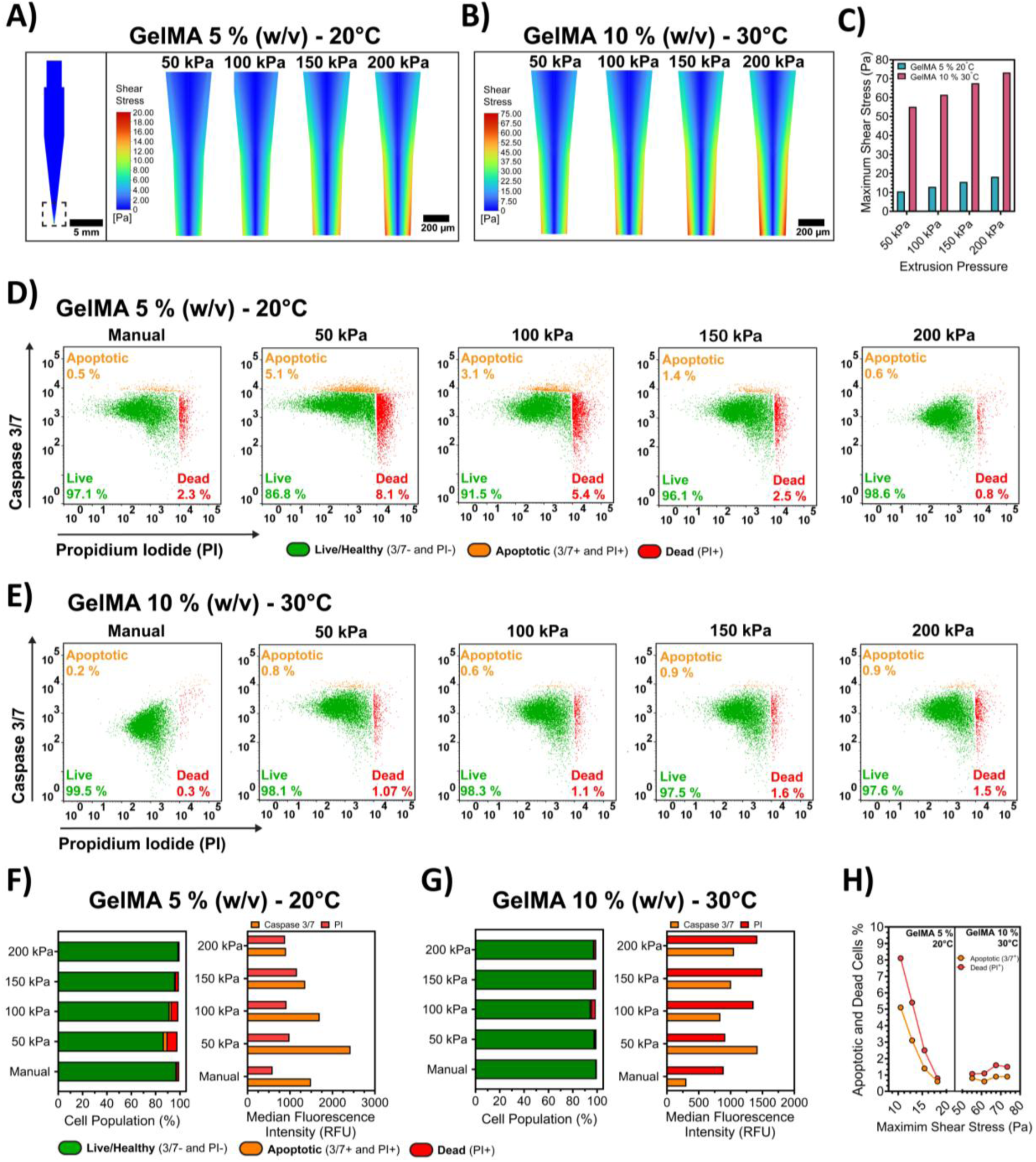
The immediate effect of bioprinting-induced shear stress before crosslinking on HUVECs and MCF-10A’s. A-B) Computational fluid dynamic (CFD) simulation of shear stress (Pa) contours at the outlet of 27G across a range of extrusion pressures (50-200 kPa) C) CFD simulation of maximum shear stress at outlet of 27G nozzle. All simulations utilize an axisymmetric model, and contour profiles of shear stress (A and B) are mirrored for accurate representation. D) Flow cytometry analysis of Caspase 3/7 and Propidium iodide (PI) in GelMA 5 % (w/v) at 20 °C comparing manually prepared precursor solution and bioprinted at different extrusion pressures (kPa). E) Flow cytometry analysis of Caspase 3/7 and Propidium iodide (PI) in GelMA 10 % (w/v) at 30 °C. F-G) Quantitative analysis of cell viability and apoptosis in GelMA 5 % (w/v) at 20 °C and GelMA 10 % (w/v) at 30 °C by quadrant gating and median fluorescence intensity (RFU) of Caspase 3/7 and PI. H) Effect of shear stress (Pa) on apoptotic dead cell (PI^+^) population in GelMA 5 % (w/v) at 20 °C and GelMA 10 % (w/v) at 30 °C.

Day-1 responses were assessed following printing and crosslinking to determine whether cells exhibited a delayed apoptotic response. GelMA 5 % (w/v) was utilized based on its previously observed increase in immediate apoptotic response compared with GelMA 10 % (w/v) (Figure 3). Manual free-swelling droplet controls were prepared at 37 °C and 20 °C, while printed groups were generated at 50, 100 and 200 kPa. Representative maximum-projection images demonstrated a reduction in live-cell coverage with increasing extrusion pressure, particularly in the 100 and 200 kPa groups (Figure 4A). Metabolic activity and DNA content normalised to hydrogel wet weight were comparable between the manual 37 °C, manual 20 °C and 50 kPa groups (Figure 4B i–ii). In comparison, extrusion at 100 and 200 kPa resulted in reduced metabolic activity and DNA content after 1 day. Quantification of stained area coverage also demonstrated greater live-cell and nuclei coverage in the manual 37 °C, manual 20 °C and 50 kPa groups compared with the 100 and 200 kPa groups (Figure 4B iii).

**Figure 4:**
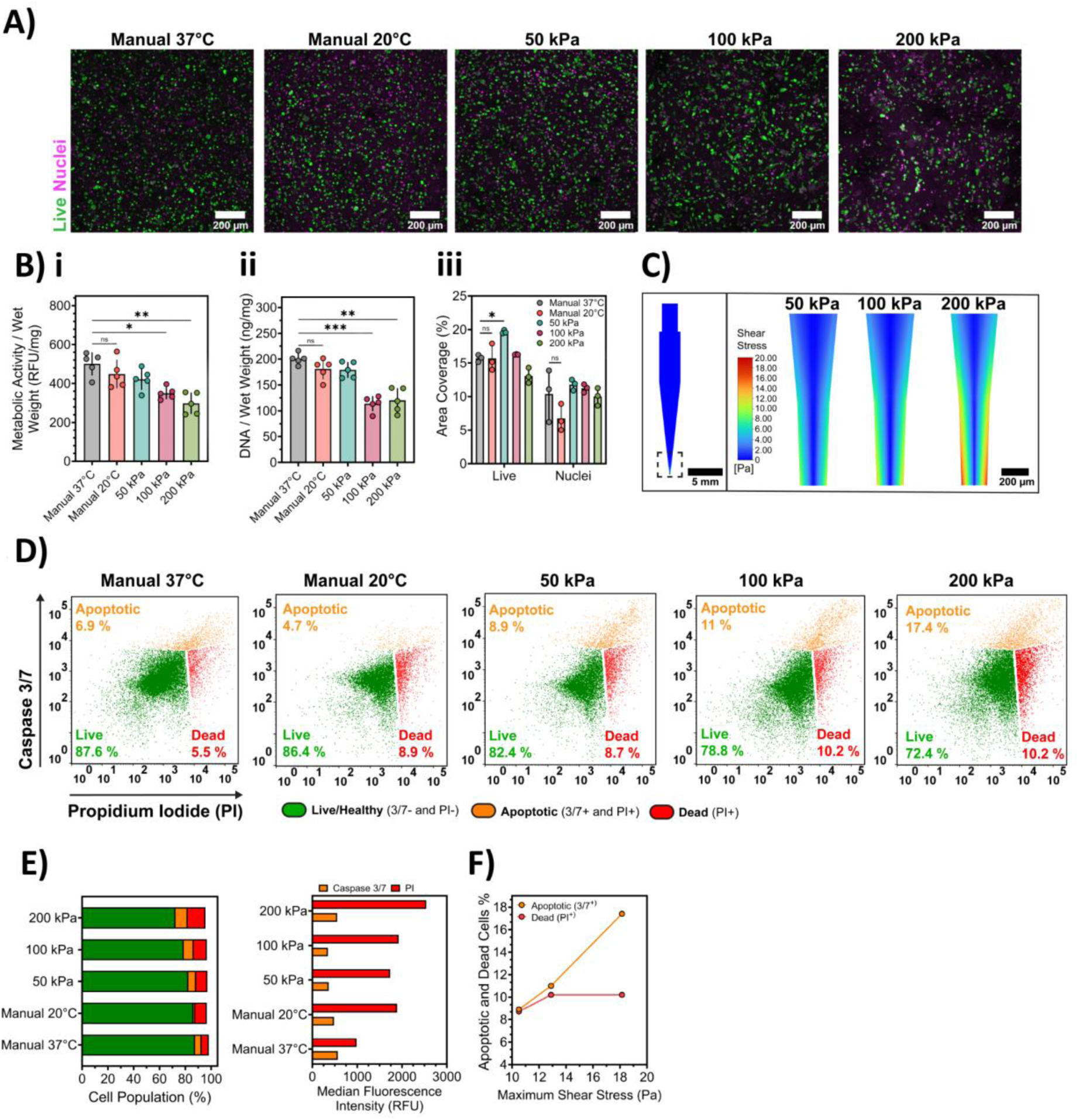
The impact of bioprinting-induced shear stress on encapsulated HUVECs and MCF- 10A’s in 5 % (w/v) GelMA after 1 day of culture. A) Representative maximum projections - cell viability of encapsulated HUVECs and MCF-10A’s in 5 % (w/v) GelMA at day 1. B) i) PrestoBlue metabolic activity (RFU) normalized to wet weight (mg); ii) DNA content (ng) normalized to wet weight (mg) (n = 5); and iii) Stained area coverage (%) of live cells (Fluorescein Diacetate – FDA, green) and nuclei (Hoechst 33342, magenta) quantified in CellProfiler, average from three ROIs per sample. C) Computational fluid dynamic (CFD) simulation of shear stress (Pa) contours at the outlet of 27G across a range of extrusion pressures (50,100 and 200 kPa). D) Flow cytometry analysis of digested hydrogels (5 % w/v GelMA) at day 1 using Caspase 3/7 and Propidium Iodide (PI). E) Quantitative analysis of GelMA 5 % (w/v) cell populations by quadrant gating and median fluorescence intensity (RFU) of caspase 3/7 and PI. F) CFD simulation of maximum shear stress at the outlet of 27G. Mean values with error bands representing standard deviation (SD), statistical analysis was performed for B) i-iii) using one-way ANOVA with Tukey post-hoc test (*P < 0.05, **P < 0.01, ***P < 0.001). All simulations utilize an axisymmetric model, and contour profiles of shear stress (C) are mirrored for accurate representation.

CFD simulations of GelMA 5 % (w/v) at 20 °C predicted maximum shear stresses of approximately 10-18 Pa at the outlet of the 27G nozzle, with shear stress increasing as extrusion pressure increased from 50 to 200 kPa (Figure 4C). The shear-stress contour profiles demonstrated an increase in the magnitude of shear stress near the nozzle wall across the extrusion-pressure range. Flow cytometry analysis of cells recovered from GelMA 5 % (w/v) hydrogels after 1 day demonstrated comparable live-cell populations between the manual 37 °C and 20 °C controls, with 87.6 % and 86.4 % live cells, respectively (Figure 4D–E). However, the manual 37 °C control had a higher apoptotic cell population (6.9 %, Caspase 3/7+/PI+), whereas the manual 20 °C control had a higher dead-cell population (8.9 %, PI+). Following bioprinting, the live-cell population decreased with increasing extrusion pressure, from 82.4 % at 50 kPa to 78.8 % at 100 kPa and 72.4 % at 200 kPa.

The apoptotic cell population increased with extrusion pressure, from 4.7 % in the manual 20 °C control to 8.9 %, 11.1 % and 17.4 % at 50, 100 and 200 kPa, respectively (Figure 4D–E). In comparison, the percentage of dead cells remained more comparable between the manual 20 °C control and the bioprinted groups, ranging from 8.7 % to 10.2 %. Median Caspase 3/7 fluorescence also increased with extrusion pressure, with the highest fluorescence observed at 200 kPa. PI fluorescence increased across the bioprinted groups, alongside the increase in the Caspase 3/7+/PI+ apoptotic population. When compared with the CFD-predicted maximum shear stress, the apoptotic population increased from 8.9 % at approximately 10 Pa to 17.4 % at approximately 18 Pa (Figure 4F). In comparison, the dead-cell population increased from 8.7 % to 10.2 % across the same shear-stress range and remained similar between the 100 and 200 kPa groups.

## Discussion

Extrusion-based bioprinting is a popular biofabrication technique used in tissue engineering, allowing for the precise deposition of bioinks and biomaterial-cell suspensions through a nozzle. Recent advancements have focused on the incorporation of computational fluid dynamics (CFD) to improve print optimization, mixing methods and study of shear stress [29,43,48,49]. However, the effectiveness of simulations depends on accurate rheological modelling of the bioinks used. This study established a proof-of-concept workflow linking GelMA rheology, non-Newtonian CFD, micro-PIV validation, and acute co-culture responses during extrusion-based bioprinting.

Utilizing rheological data of bioinks to predict their behavior during bioprinting is a well-established analytical method that provides valuable insights [2,10,15]. Non-Newtonian regression models, such as Power-law, Herschel-Bulkley, Cross, and Carreau, are commonly used to identify flow behavior under varying shear rates [38,50]. However, the appropriate application of these models is critical and depends on the specific rheological data. In our study, we demonstrate the fit of rheological data for GelMA 5 % (w/v) at 20 °C and 10 % at 30 °C using these non-Newtonian models, highlighting the importance of model selection in accurately representing bioink behavior across different shear rate ranges for computational fluid dynamics of bioprinting-associated shear stress modelling. GelMA demonstrated known temperature-dependent viscosity and shear-thinning behavior in all concentrations (Figure 1A, i) [51].

In comparison of non-Newtonian regression models for GelMA 5 % and 10 % (w/v), the Carreau model demonstrated superior performance across a shear rate range of 0.1-1000 s^-1^ with experimental data (R^2^ values 0.949-1.0). In contrast, the Power Law, Herschel-Bulkley and Cross models are known to underestimate viscosity at higher shear rates (1-1000 s^-1^), despite showing similar flow behavior indices (Figure 1A i-ii, Supplementary Data Table S1, Figure S8) [43,50,52]. Simulated shear rates using the Carreau model for GelMA 5 % (w/v) at 20 °C ranged from 830 to 1,435 s⁻¹, while GelMA 10 % at 30 °C, ranged from 5,854 to 7,784 s⁻¹, using extrusion pressure of 50 to 200 kPa. These wide ranges of shear rates highlight the importance of selecting a model that accurately captures the materials flow behavior. The Power Law and Herschel-Bulkley models, which tend to underestimate viscosity at higher shear rates, may not accurately represent the material behavior under these printing conditions.

CFD simulations using ANSYS® Fluent revealed minimal differences in shear stress and velocity profiles between regression models at 40 kPa extrusion pressure for both GelMA concentrations (Figure 1B-C). However, the Carreau model excelled in capturing complex rheological behavior of biomaterials like GelMA for support-bath printing, gelatin-carboxymethyl cellulose, and alginate blends for extrusion printing [43,50]. To effectively rely on data obtained from computational fluid dynamics (CFD) simulations, robust validation is essential. Previous studies have shown the effectiveness of CFD simulations in investigating the influence of nozzle geometries, screw-based dispenser systems, and predicting shear stress on HUVECs in 3-4 % (w/v) GelMA or HepG2 viability [30,31,53]. However, many of these studies lack validation of their simulations or a robust workflow to emphasize this crucial step. In the current study, micro-PIV was employed to validate the predictions of the Carreau material model. Micro-PIV measurements of water and GelMA 10 % (w/v) at 30 °C (Figure 4.2) aligned well, validating the selected model. Although CFD has been increasingly applied to extrusion bioprinting, many studies focus primarily on predicting shear stress or printability without experimentally validating model outputs or integrating rheological characterization into the simulation workflow. By combining rheological modelling, CFD simulations and experimental micro-PIV validation, the present workflow provides a framework for predicting extrusion-associated shear stress before biological experimentation. Such an approach may reduce empirical optimization during bioink development while improving reproducibility across different material formulations and printing conditions [54].

After validating the CFD simulations, initial cell studies investigated the immediate impact of bioprinting-associated shear stress on cell viability prior to crosslinking and Day-1 response post crosslinking. These studies were performed without crosslinking and examined the effects of extrusion pressures ranging from 50 to 200 kPa. For GelMA 5 % (w/v) at 20 °C, a higher proportion of cells expressed apoptosis marker (Caspase 3/7^+^/PI^+^) across all experimental printing groups ranging from 60.6 % to 5.1 %, with 50 kPa demonstrating greater apoptotic expression (Figure 3A). In contrast, GelMA 10 % (w/v) at 30 °C groups showed negligible difference of cells expressing apoptosis (0.2-0.9 %, Caspase 3/7^+^/PI^+^) (Figure 3E).

To effectively bioprint GelMA, printing temperature is modulated and set depending on the polymer concentration. In this current study, non-crosslinked GelMA 5 % (w/v) at 20 °C had increased presence of apoptotic cells in printed groups compared to manual groups. Interestingly, within printed groups apoptosis decreased with extrusion pressure, which may be attributed to an increase in mass flow rate and reduced exposure duration to shear stress (Figure 3A). In contrast, GelMA 10 % (w/v) at 30 °C demonstrated limited differences between printing conditions and manual control groups (Figure 3E) [85, 144]. Cells printed and crosslinked in 5 % (w/v) GelMA at 20 °C exhibited clear differences in apoptotic and dead cell markers after 1 day compared to immediate post-printing (Figure 3D and 4D).

Apoptosis is triggered by through intrinsic cellular damage or external death receptor signaling, resulting in activation of caspase cascade. Caspases are broadly classified into initiator caspases (caspase-8, -9, and -10), which activate executioner caspases (caspase-3, -6, and -7) responsible for cellular dismantling [55]. Activation of executioner caspases, particularly caspase-3 and caspase-7, has traditionally been considered an irreversible commitment to apoptosis. [56,57]. Although the activation executioner caspases such as caspase-3 and -7 have been considered irreversible with cells However, recent studies suggest cells are capable of recovering from early-stage apoptosis through a process known as anastasis, provided initiating cellular stress is limited [56–58]. This suggests that early activation of caspase-3/7 following extrusion may represent an acute stress response rather than irreversible cell death.

The flow cytometry analysis does not reveal distinct populations of live/healthy, apoptotic, and dead cells (Figure 3D-E and 4D). Rather than representing discrete biological states, apoptosis is a dynamic and continuous process in which cells progressively transition from healthy to apoptotic and ultimately secondary necrotic states. Healthy cells may express low levels of Caspase 3/7 during normal homeostatic functions, and progression from apoptosis to secondary necrosis involves overlapping cellular processes [56]. Unlike specific antibody labelling, which often results in more discrete populations, caspase activity assays can detect a range of activation levels.

Differences were observed between manual groups (20 °C and 37 °C) and the 50 kPa extrusion groups’ live populations (82.4% to 87.6 %), suggesting that extrusion pressures below 50 kPa may have little impact on cellular function compared to extrusions pressures ranging from 50 to 200 kPa (Figure 4D Collectively, these findings indicate that both printing temperature and extrusion pressure influence the acute cellular response to bioprinting by altering the environment experienced by cells[59]. This response was reflected by reduced metabolic activity and DNA content, which became more pronounced with increasing extrusion pressure (Figure 4B i-ii) [60,61]. Although extrusion bioprinting is commonly associated with elevated shear stress, manual pipetting may result in comparable cellular viability outcomes. This similarity can be attributed to the potentially higher and more variable flow rates during manual pipetting, potentially exposing cells to similar or even higher levels of shear stress during rapid manual dispensing.

The CFD simulations predicted maximum shear stresses of 10-18 Pa for GelMA 5 % (w/v) at 20 °C and 55-73 Pa for GelMA 10 % at 30 °C (Figure 3A-B). . These differences are consistent with distinct rheological behaviour of the two bioinks. Although GelMA 10% (w/v) at 30 °C exhibited more Newtonian-like behaviour, its viscosity remained substantially higher than that of 5% (w/v) GelMA under printing conditions, resulting in greater predicted wall shear stresses. In contrast, GelMA 5% (w/v) at 20 °C exhibited stronger shear-thinning behaviour, reducing its apparent viscosity during extrusion and consequently lowering the predicted shear stress. (Table S4.1, Figure S4.8 – Supplementary Data). Previous studies using CFD simulations to model bioprinting-associated shear stress have reported values, ranging from 5 to 10,000 Pa, reflecting differences in bionk rheology, printing parameters, nozzle geometry and cell type [19–21,28,29,31,49,60].[9,10]. Many of these studies employ substantially more viscous bioinks, such as alginate (0.5-1 % w/v), which exhibit viscosities approximately 10-10,000-fold higher than the GelMA formulations used in this study. Consequently, substantially higher simulated shear stresses would be expected under otherwise comparable printing conditions.

[20,21,25,28,29]. A comparable study using GelMA 3-5 % (w/v) reported simulated shear stress values of 10-300 Pa while maintaining cell viability ≥ 70 % over 7 days using a constant feed rate of 100 µl min^-1^ [31]. Under comparable GelMA concentrations, our study demonstrated day 1 cell viabilities of 72.4-87.6 % (Figure 4). Differences between studies may reflect several experimental variables, including the use of a HUVEC-MCF10A co-culture rather than a monoculture, differences in GelMA source, and distinct viability assessment time points. Here, we demonstrated a validated workflow integrating rheological characterization, CFD modelling and experimental validation to predict acute bioprinting-associated shear stress responses in multicellular constructs. As extrusion bioprinting continues to advance towards increasingly complex bioinks and co-culture systems, predictive modelling frameworks that incorporate experimentally derived material properties may reduce iterative optimization while improving reproducibility during bioink development. Although the present study focused on acute and day-1 responses, the workflow is readily adaptable to alternative bioinks, nozzle geometries and cell types through incorporation of material-specific rheological inputs. Future work should therefore extend this framework to longer culture periods, additional biomaterial systems and more mechanically sensitive cell populations. Ultimately, integrating this workflow into bioprinting platforms could enable real-time optimization of printing parameters to minimize cell damage while maintaining print fidelity.

## Methods

### GelMA

Sterile lyophilized Gelatin Methacryloyl (GelMA; porcine skin, Type A, 80 % degree of functionalization) was obtained from Gelomics Pty Ltd (Brisbane, Australia). GelMA stock solutions (22.5 % w/v) were prepared in phosphate-buffered saline (PBS) (Life Technologies, Thermofisher, Waltham, MA, United States) overnight at 37 °C in a rotating oven. Lithium phenyl-2,4,6-trimethylbenzoylphosphinate (LAP) (Merck, MA, USA) photoinitiator was dissolved in PBS to obtain a 3 % (w/v) stock solution and sterilized using 0.22 µm membrane filter (Merck). Final GelMA precursor solutions were prepared at 5 % and 10 % (w/v) with 0.15 % (w/v) LAP photoinitator in PBS and protected from light in a 37 °C water bath.

### Rheology Preparation

The viscosity profile of each GelMA precursor solution was determined using a MCR302 rheometer (Anton Paar, Graz, Austria) with a double-gap (DG26.7) geometry using a logarithmic shear rate ramp (0.1 – 1000 s^-1^). The DG26.7 geometry was selected instead of traditional cone plate geometry due to GelMA’s low viscosity at testing temperatures. Final GelMA precursor solutions were prepared fresh and kept at 37 °C in a dry oven for 15 min, then allowed to cool to room temperature (25 °C) for 30 min before loading in DG geometry. Each replicate (n = 3) required 3-4.5 ml of precursor solution, each aliquot was allowed to equilibrate at test temperature (20 and 30 °C) for 15 min before test commencement.

Based on the rheological data obtained, regression models were applied to define input parameters for non-Newtonian simulations, with each equation based on ANSYS® Fluent models [62]. The regression models used included the Power Law (Eq. 1), Herschel-Bulkley (Eq. 2), Cross (Eq. 3) and Carreau (Eq.4).

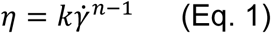

Where *k [Pa·s]* is the consistency index, *γ̇* is the shear rate (s^-1^) and *n* is the power law index.

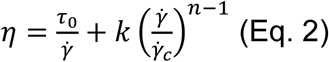

ANSYS® Herschel-Bulkley model combines Bingham plastic and power-law models for the determination of fluid viscosity, where *τ*_0_ is yield stress *[Pa], γ̇* is the shear rate (s^-1^), *k [Pa·s]* is the consistency index, *γ̇_c_* is the critical shear rate (s^-1^) and *n* is the power law index.

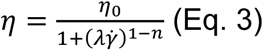

Where *η*_0_ *[Pa·s]* is the zero-shear-rate viscosity, *λ* is natural time (s), *γ̇* is the shear rate (s^-1^) and *n* is the power law index.

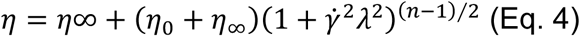

### Design of Experiment Mass Flow Rate

A Design of Experiment (DoE) approach was utilized to model the mass flow rate of GelMA (5 % and 10 %, w/v) and water within the printing pressure range (20 – 200 kPa) using DoE software MODDE® (Version 13, Umetrics™, Sartorius Stedim Biotech, Aubagne, France). A level design was used, with 10 design runs, 2 center points and 4 replicates, resulting in 65 total runs. A BIO X (CELLINK, Gothenburg, Sweden) 3D Extrusion Bioprinter was used for all printing with the temperature-controlled printhead. All materials were inserted into 3 ml syringes (Nordson, Westlake, Ohio, United States) and loaded into the printhead for 20 min at a set temperature (20 °C – 5 % w/v GelMA, 25 °C – water, 30 °C – 10 % w/v GelMA). First, empty 2 ml Micro Tubes (Sarstedt AG, Nümbrecht, Germany) were weighed using Cubis II Semi Microbalance (Sartorius, Göttingen, Germany), a custom G-code was utilized to extrude material for 1 s or 5 s into the designated tube for each replicate/pressure per material. Subsequently, each tube was weighed, and the mass of the empty tube was subtracted to determine the total extruded mass (mg). This value was then divided by the corresponding extrusion time to calculate the mass flow rate (mg/s).

### Micro-computed tomography

To accurately model the 3 ml syringe barrel and 27G (200µm) nozzle (Nordson, Westlake, Ohio, United States) micro-computed tomography (µ-CT) was performed to obtain measurements using µCT50 (Scanco Medical AG, Wangen-Brüttisellen, Switzerland) in combination with supplied dimensions from the manufacturer. To reduce scan time and software processing the bottom ¼ of the syringe barrel and the 27G nozzle were scanned using a voxel size of 14.6 µm, energy of 55 kVp and current of 145 μA. Measurements were obtained through 3D reconstruction and cross-sections processed using Image J (National Institutes of Health, Bethesda, MA, USA) and SOLIDWORKS (Waltham, MA, USA).

### Computational Fluid Dynamics

Computational fluid dynamic (CFD) analysis was performed using ANSYS® Academic Research Fluent 19.2 (ANSYS, Canonsburg, PA, USA). All simulations were performed on Dell Latitude 7420 using an Intel Core i7-1185G7 with 16 GB RAM running a Windows 10 (64-bit) operating system. The reconstructed syringe barrel and 27G nozzle model were created in Fluent DesignModeler as a 2D axisymmetric model to minimize computational processing requirements (Figure 1S, Supplementary Data). Next, a non-uniform mesh was applied to capture the geometry using standard CFD preferences adapting the element, face and edge sizing (≥ 1-0.015 mm). The mesh size was adaptively refined as the geometry tapered to ensure flow characteristics were represented. A viscous laminar model was applied using a steady-state pressure-based solver, absolute velocity and axisymmetric 2D space. GelMA 5 % (w/v) at 20 °C and GelMA 10 % at 30 °C were created as new fluid materials in ANSYS® Fluent, water was utilized from the existing Fluent database. For the material model of GelMA materials, the density was determined to be 1031.7 kg/m^3^ using Ultrapyc 5000 (Anton Paar). Non-Newtonian models (Power-Law, Herschel-Bulkley, Cross and Carreau) were selected for each GelMA material and model parameters were input from the regression modelling of each equation (Table S1, Supplementary Data). Based on the DoE prediction model, and given mass flow is consistent throughout an enclosed conduit, the mass flow at simulated extrusion pressures was used to calculate the inlet velocity (mm/s) using the following equation (Eq. 5):

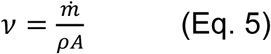

Where *ν* (m/s) is the velocity, *ṁ* is the mass flow rate (kg/s), *ρ* is the density (kg/ m^3^) and *A* the inlet area (m^2^).

In Fluent the outlet was set as an Outflow with a flow rate weighting of 1, the solution was initialized using Hybrid Initialization with Coupled Scheme, Least Squares Cell Based Gradient, Second Order Pressure and Second Order Upwind Momentum. An independent mesh convergence study (Figure S2, Supplementary Data) was performed using a series of 8 meshes (M1-M8), progressively refining element sizes in each iteration from the coarsest mesh (M1) to the finest mesh (M8). The Fluent convergence study was performed with GelMA 10 % (w/v) – 30 °C using the Carreau model at 40 kPa extrusion pressure. All data exported from the axisymmetric model in ANSYS® Fluent was mirrored for enhanced representation of the nozzle profile.

### Micro-Particle Image Velocimetry

To validate the CFD simulation velocity profiles of water and GelMA in the nozzle geometry, Micro-Particle image velocimetry (micro-PIV) was conducted. A 3D printed phantom model of the 27G nozzle was designed with SOLIDWORKS 2023 (Dassault Systems, France) (Figure S6, Supplementary Data). The model STL file was initially prepared using ASIGA Composer Software (Version 1,3, ASIGA), and later fabricated with Asiga Max X27 DLP 3D printer (ASIGA, Australia) with Mooin TechCLear resin (DMG Digital Enterprises SE, Germany). Postprocessing of the phantom included a 5 min wash in isopropanol (IPA, Sigma-Aldrich, Australia), and a 2 hr UV curing time. The micro-PIV setup utilized a syringe pump (Legato® 100 Syringe Pump, KD Scientific, USA) with a 10 ml syringe attached, connecting to the 3D printed model via silicone tubing. Volumetric flow rates for each material (water and GelMA) were set on the peristaltic pump based on previously calculated mass flow rates. Micro-PIV utilized 1-20 µm sized fluorescent particles (PMMARhB-Frak-Particles, DANTEC, Germany) and were excited at 560 nm and emitted light at 584 nm, which was captured by the fluorescence filter of the microscope. The fluorescent particles were added to a mixture of water and GelMA. A 10.56 % (w/v) GelMA stock concentration was prepared, 5 % (w/v) aqueous suspension of particles was added to GelMA to achieve a final concentration of 10 % (w/v). The Micro-PIV system (Dantec Dynamics A/S variant, Denmark) contains a DualPower laser (100/200 Hz, Dantec Lasers) initiated by a synchronizer connected to the high-speed camera-equipped microscope. A 2.5x magnification objective was used to visualize the flowing particles in the printed nozzle phantom. Three image sets, each containing 10 pairs of images, were captured at 10 Hz defined trigger rate and 4160 µs intervals between pulses. Captured images were processed in MATLAB (MathWorks, Natick, MA, USA) software with the PIVLab plugin [63]. Each image underwent background reduction and masking following a multiphase cross-correlation algorithm to obtain a vector map of particle velocities within the model. This data was then exported from MATLAB and correlated with velocities obtained in CFD simulations for water and 10 % (w/v) GelMA.

### Cell culture

HUVECs were isolated as previously described [64] from one donor and used at passages 3-4, cultured in endothelial basal medium (EBM™-2, Capsugel®, Lonza, Sydney, Australia) and supplemented with EGM™-2 SingleQuots supplement (Capsugel®, Lonza). The MCF-10A human breast epithelial line were kindly gifted by Professor Derek Richard (Queensland University of Technology) and were used at passages 6-7 and cultured in DMEM/F12 GlutaMAX™-1 supplemented with 5 % (v/v) horse serum, 1 % (v/v) penicillin-streptomycin (both Gibco™, Thermofisher, Brisbane, Australia), 20 ng/ml human epidermal growth factor (hEGF, Miltenyi Biotec, Sydney, Australia), 0.5 µg/ml hydrocortisone, 10 µg/ml human insulin and 100 ng/ml cholera toxin (all Merck) and maintained as described previously [65]. All cells were seeded in T175 flasks (Nunc™, Thermofisher), cultured to 70-80 % confluency at 37 °C in a humidified incubator containing 5 % CO_2_, cells were monitored daily, and media change was performed every 2-3 days.

### 3D Cell Culture and Bioprinting

For hydrogel co-culture encapsulation, HUVECs and MCF-10A cells were first washed with PBS before detachment with 0.25 % (v/v) trypsin EDTA (Gibco™) and counted with 0.4 % (v/v) trypan blue solution (Gibco™) using an automated cell counter (Invitrogen, Thermofisher). In all experiments, HUVECs were encapsulated at a density of 6 × 10^6^ cells/ml and MCF-10A at a density of 6 × 10^5^ cells/ml. HUVECs and MCF-10A’s were encapsulated into 5 % (w/v) and 10 % GelMA to assess the direct effect of bioprinting before crosslinking. Each cell-laden bioink was loaded into a temperature-controlled printhead and allowed to reach the desired printing temperature of 20 °C (5 % w/v GelMA) or 30 °C (10 % w/v GelMA) for 30 min. The bioinks were then printed with the BioX using 27G conical nozzle into 48-well plates (Thermofisher) using custom GCODE based on mass flow data previously collected for each material at desired extrusion pressures (50, 100, 150 and 200 kPa) to achieve a 100 µl cell suspension in triplicates. Manual control groups for 5 % and 10 % w/v GelMA were prepared at the same cell density and temperature using Finntip P200 pipette (Thermofisher). Next HUVECs and MCF-10A’s were encapsulated in same density previously used in 5 % (w/v) GelMA and bioprinted into 48-well plate to produce 20 µl free-swelling droplets for each extrusion pressure (50, 100 and 200 kPa) based on mass flow prediction data. Manual hydrogels were prepared at identical density to printed at 20 °C and 37 °C, all hydrogels were crosslinked for 45 s using the LunaCrosslinker™ Visible Light Crosslinking System (Gelomics Pty Ltd) and submerged in EGM-2 cell culture media, incubated at 37 °C with 5% CO_2_.

### Flow Cytometry

The cell suspension and hydrogels were digested in 1 mg/ml Collagenase Type I (Baoding Faithful Industry Co, Hebei, China) for 10 mins at 37 °C. After digestion, Collagenase was neutralized using EGM-2 media, samples of each group (n=5) were pooled and then centrifuged at 1000 rpm for 5 mins. The recovered cells were then resuspended in 1 ml of FACS Buffer (Miltenyi Biotec). Fluorescence Minus One (FMO) control was established using manually pipetted groups and cells from culture flasks for each flow cytometry experiment and included unstained, propidium iodide and caspase 3/7 for correct gating of fluorophores. To ensure positive staining in the controls, two methods were employed. For apoptosis induction, cells attached to tissue culture flasks were treated with 0.5 mM H_2_O_2_ added to the cell culture media and incubated overnight at 37 °C with 5 % CO_2_, as described previously (Figure S4, Supplementary Data) [66]. To assess the positive dead cell population, cells were first detached and centrifuged to remove the supernatant. Then, 1 ml of 70 % (v/v) filtered ethanol was added to the cell pellet and incubated at room temperature for 30 minutes. After this incubation period, the ethanol was removed by centrifugation, and the cell pellet was resuspended in 1 ml of FACS buffer. Briefly, cell suspensions were incubated with a final concentration of 0.5 µM CellEvent™ Caspase-3/7 (Thermofisher) for 45 mins at 37 °C and stained with 5 µg/ml of PI for 2 mins prior to each sample being run. Each sample was filtered through 35 µm cell strainer 5 ml Falcon® test tube (Corning, NY, USA).

Cell acquisition was performed using BD FACSCelesta™ (BD, NJ, USA) flow cytometer and analyzed using FlowJo software version 10.10 (FlowJo LLC, BD, NJ, USA). Optimal voltage settings were determined using FMO controls and a representative sample. Initial gating strategy identified main cell population (P1) and single cells. Compensation was performed by the collection of 5 − 10 × 10^4^ single cell events for each FMO control. Subsequently, experimental samples were analysed and collected 2.0 − 3.0 × 10^4^ single events per sample. Negative population thresholds for Caspase 3/7 and PI were established using unstained controls, with gates set to include 98 % of the unstained cells in each channel, ensuring standardized analysis across all experimental samples.

### Cell Viability

The viability of cell-laden hydrogels was assessed using Fluorescein Diacetate (FDA, final concentration of 10 µg/ml, Thermofisher) in PBS and Hoeschst 33342 (final concentration of 10 µg/ml, Thermofisher). Samples were imaged using Olympus FV4000 confocal microscope (Evident, Olympus Life Science, Tokyo, Japan) at 10x magnification for volumetric z-stacks with an average depth of 400 µm and 1 µm slices using resonant scanner with a resolution of 1024x1024. Each sample had three regions of interest (ROI) imaged randomly followed by quantification using Image J and CellProfiler version 4.2.7 (Broad Institute, MA, USA).

### Presto Blue Metabolic Assay

Metabolic activity of D1 co-culture samples was assessed using PrestoBlue Cell Viability Reagent (Invitrogen™, Thermofisher) according to the manufacturer’s recommendation. Briefly, a 1:10 ratio (PrestoBlue: cell culture media) was prepared and 500 μl of reagent solution was added per well and incubated for 45 min at 37 °C. Triplicates of 100 μl from each sample were aliquoted into a 96-well plate, and relative fluorescent units (RFU) was read at 590 nm using a microplate reader (BMG LABTECH, Ortenberg, Germany) at two different gains (900 and 1200). Data was normalized to wet weight of each sample to account for variations in printing and manual groups.

### DNA Quantification

Cell-laden hydrogels were weighed to obtain weight wet, then transferred to 2 mL centrifuge tubes and frozen at −80 °C. Samples were then digested overnight at 65 °C with 0.5 mg/ml Proteinase K (Invitrogen™, Thermofisher) in phosphate-buffered EDTA (PBE, pH 7.1). DNA content was quantified using Quant-iT™ PicoGreen^®^ (Life Technologies) assay. DNA standards were prepared with concentrations ranging from 1000 ng/ml to 31.25 ng/ml. Samples were diluted 1:5 in PBE and transferred in triplicate to a 96-well plate. PicoGreen^®^ dye (100 μl) was added to each well, and the well-plate was incubated for 5 min at room temperature protected from light. Fluorescence was measured at an excitation wavelength of 480 nm and emission wavelength of 520 nm, using a CLARIOstar^®^ Plus spectrophotometer (BMG Labtech, Mornington, Australia). The DNA content of sample digests was determined by comparison of sample fluorescence to DNA standard curve and normalized to hydrogel weight.

## Statistical and data analysis

Statistical analysis was performed using GraphPad Prism version 10.0. All data shown as mean ± standard deviation (SD), n = 5 technical replicates and described in each figure caption unless stated otherwise. Comparisons between two groups were analyzed using Student’s t-test with Welch’s correction. Comparison between multiple groups utilized One-Way ANOVA, followed by Tukey or Bonferroni multiple comparison tests. A p-value of less than 0.05 was considered statistically significant with asterisks denoting statistical value (*P < 0.05; **P < 0.01; ***P < 0.001).

## Acknowledgments

All data reported was supported by Queensland University Technology’s Central Analytical Research Facility.

## Author Contributions

J.W.D., A.W and T.J.K conceptualized the studies and methodologies; J.W.D., A.W and J.A.C. performed experimentation and analysis; J.W.D and A.W drafted the original manuscript; L.J.B., C.M. and T.J.K. provided resources and funding; J.W.D., A.W., J.A.C., C.M., L.J.B and T.J.K. all authors have revised, read and agreed to the published version of the manuscript.

