## Supplementary data for "Modelling and measuring effects of shear stress in extrusion bioprinting of endothelial- epithelial cell co-cultures" (from another version of this preprint: version 1, 2026-09-03)

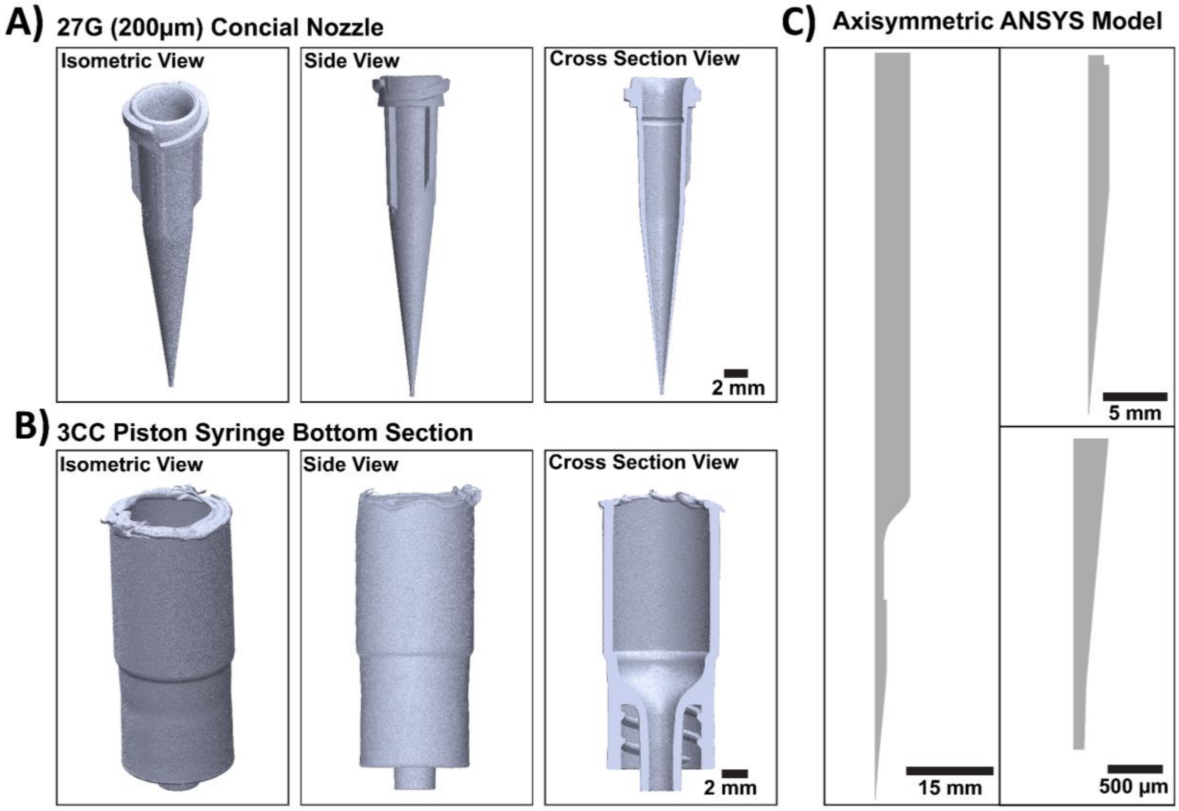


**Figure S1: 3D volume rendered from micro-computed tomography (µ-CT) and generated axisymmetric geometry model.** A) 3D render of 27G conical nozzle in SOLIDWORKS B) 3D render of the bottom section of 3cc piston syringe in SOLIDWORKS C) Axisymmetric geometry of 27G conical nozzle and 3cc piston syringe, the model generated in DesignModeler.

**Table S1: Properties of non-Newtonian model's comparison.**

| | N index | Consistency Index, K (Pa.s) | Critical Shear Rate, γ̇c ($s^{-1}$) | Natural Time, λ (s) | Yield Stress Threshold, τ0 (Pa) | Zero-shear viscosity, η0 (Pa s) | Infinite-shear viscosity, η∞ (Pa s) |
|---|---|---|---|---|---|---|---|
| **GelMA 5 % (w/v) – 20 °C** | | | | | | | |
| **Power Law** | 0.324 | 18.777 | N/A | N/A | N/A | N/A | N/A |
| **Herschel-Bulkley** | 0.324 | 5.034 | 7.011 | 0.142 | 0.000010 | N/A | N/A |
| **Cross** | 0.076 | N/A | 0.205 | 4.855 | N/A | 104.195 | N/A |
| **Carreau** | 0.323 | N/A | 24.490 | 0.040 | N/A | 0.0201 | 0.00000997 |
| **GelMA 5 % (w/v) – 37 °C** | | | | | | | |
| **Power Law** | 0.641 | 0.011 | N/A | N/A | N/A | N/A | N/A |
| **Herschel-Bulkley** | 0.640 | 0.015 | 0.366 | 2.731 | 0.00 | N/A | N/A |
| **Cross** | 0.488 | N/A | 0.100 | 10.00 | N/A | 0.0485 | N/A |
| **Carreau** | 0.190 | N/A | 0.395 | 2.529 | N/A | 0.0039 | 0.00395 |
| **GelMA 10 % (w/v) – 30 °C** | | | | | | | |
| **Power Law** | 0.910 | 0.015 | N/A | N/A | N/A | N/A | N/A |
| **Herschel-Bulkley** | 0.974 | 0.011 | 28.047 | 0.035 | 0.012 | N/A | N/A |
| **Cross** | 0.857 | N/A | 0.914 | 1.0929 | N/A | 0.0310 | N/A |
| **Carreau** | 0.529 | N/A | 0.679 | 1.471 | N/A | 0.009 | 0.009 |
| **GelMA 10 % (w/v) – 37 °C** | | | | | | | |
| **Power Law** | 1.00 | 0.015 | N/A | N/A | N/A | N/A | N/A |
| **Herschel-Bulkley** | 1.00 | 0.015 | 0.237 | 4.21 | 0.00 | N/A | N/A |
| **Cross** | 0.90 | N/A | 1.25 | 0.799 | N/A | 0.033 | N/A |
| **Carreau** | N/A | N/A | 0.395 | 2.527 | N/A | 0.014 | 0.014 |

A)

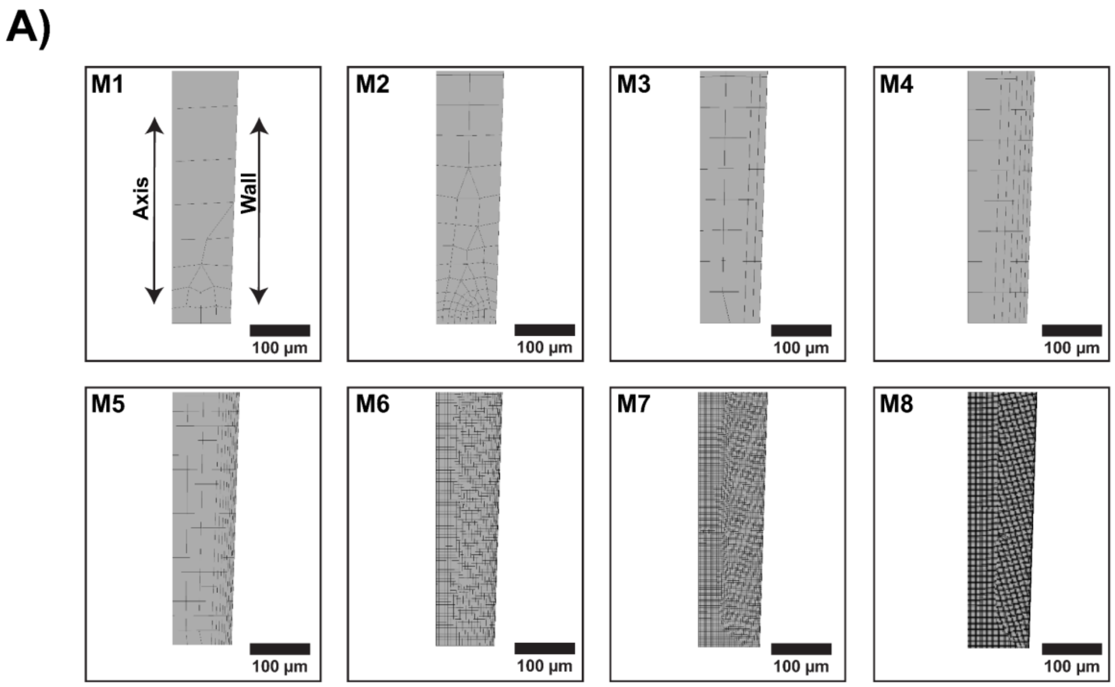


B)

**Table 2. Mesh details of each iteration (M1-8) used in the mesh convergence study.**

| | | M1 | M2 | M3 | M4 | M5 | M6 | M7 | M8 |
|---|---|---|---|---|---|---|---|---|---|
| Mesh Details | **Nodes** | 533 | 2567 | 4573 | 6007 | 14033 | 49917 | 73942 | 298908 |
| | **Elements** | 412 | 2262 | 4170 | 5608 | 13445 | 48637 | 72308 | 295583 |
| | **Mesh Method** | | | | | | | | |
| | **Element Size (mm)** | 1.00 | | | | | | | |
| | **Use Adaptive Sizing** | No | | | | | | | |
| | **Mesh Defeaturing** | Yes | | | | | | | |
| | **Defeature Size (mm)** | 0.005 | | | | | | | |
| | **Face Sizing** | None | Yes | Yes | Yes | Yes | Yes | Yes | Yes |
| | **Location** | | Face 1 | Face 1 | Face 1 | Face 1 | Face 1 | Face 1 | Face 1 |
| | **Element Size (mm)** | | 0.5 | 0.5 | 0.5 | 0.5 | 0.25 | 0.25 | 0.1 |
| | **Location** | | Face 2 | Face 2 | Face 2 | Face 2 | Face 2 | Face 2 | Face 2 |
| | **Element Size (mm)** | | 0.25 | 0.25 | 0.25 | 0.25 | 0.1 | 0.05 | 0.025 |
| | **Location** | | Face 3 | Face 3 | Face 3 | Face 3 | Face 3 | Face 3 | Face 3 |
| | **Element Size (mm)** | | 0.25 | 0.1 | 0.1 | 0.05 | 0.025 | 0.025 | 0.015 |
| | **Location** | | Face 4 | Face 4 | Face 4 | Face 4 | Face 4 | Face 4 | Face 4 |
| | **Element Size (mm)** | | 0.05 | 0.05 | 0.05 | 0.025 | 0.005 | 0.0025 | 0.001 |
| | **Inflation** | None | None | Smooth Transition | Smooth Transition | Smooth Transition | Maximum Thickness | Maximum Thickness | Maximum Thickness |
| | **Transition Ratio** | | | 0.272 (Default) | 0.272 (Default) | 0.272 (Default) | 0.02mm max | 0.02mm max | 0.02mm max |
| | **Maximum Layers** | | | 2 (Default) | 6 | 10 | 10 | 10 | 10 |
| | **Growth Rate** | | | 1.2 | 1.2 | 1.2 | 1.2 | 1.2 | 1.2 |

C)

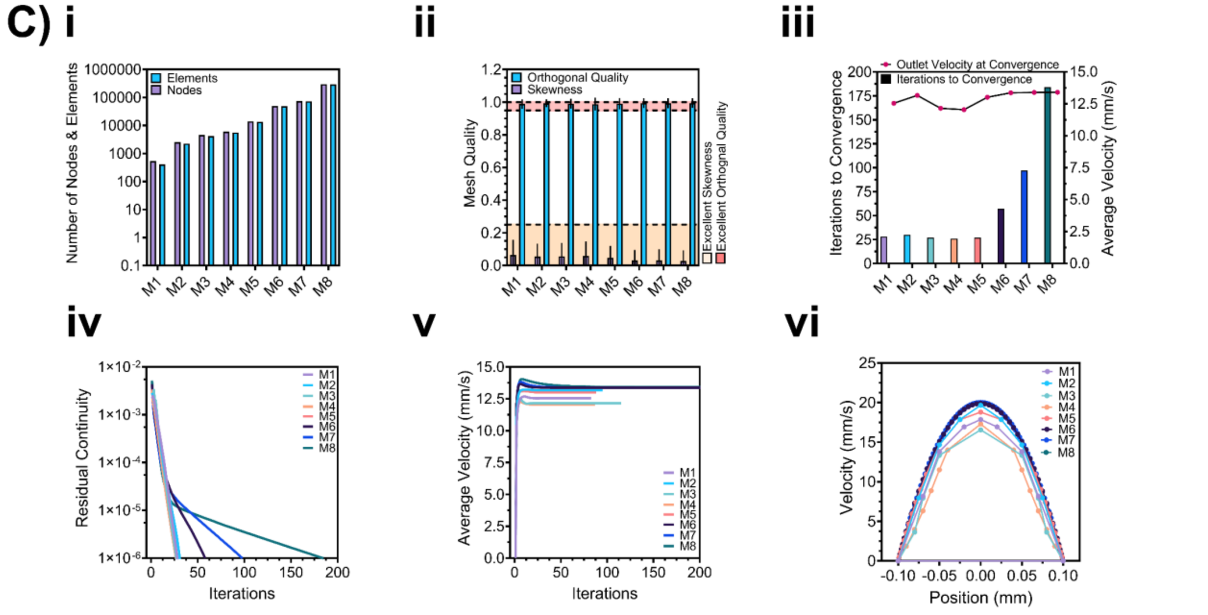


**Figure S2: Independent mesh convergence study.** A) Meshes (M1-8) utilized in convergence study at the nozzle outlet of model B) Mesh details used for each iteration M1-M8 C) Convergence results from independent study i) Total number of nodes and elements per mesh iteration ii) Quality of mesh using skewness (≤ 0.25-0, excellent skewness) and orthogonal quality (≥ 0.95-1, excellent orthogonal quality) iii) The number of iterations to reach convergence and average velocity (mm/s) at outlet per mesh refinement iv) The number of iterations required for the residual continuity to achieve convergence criteria ($1 \times 10^{-6}$) v) Average velocity (mm/s) of each mesh until converged vi) Velocity (mm/s) outlet profile of each mesh. All simulations utilize an axisymmetric model, contour profiles are mirrored for accurate representation.

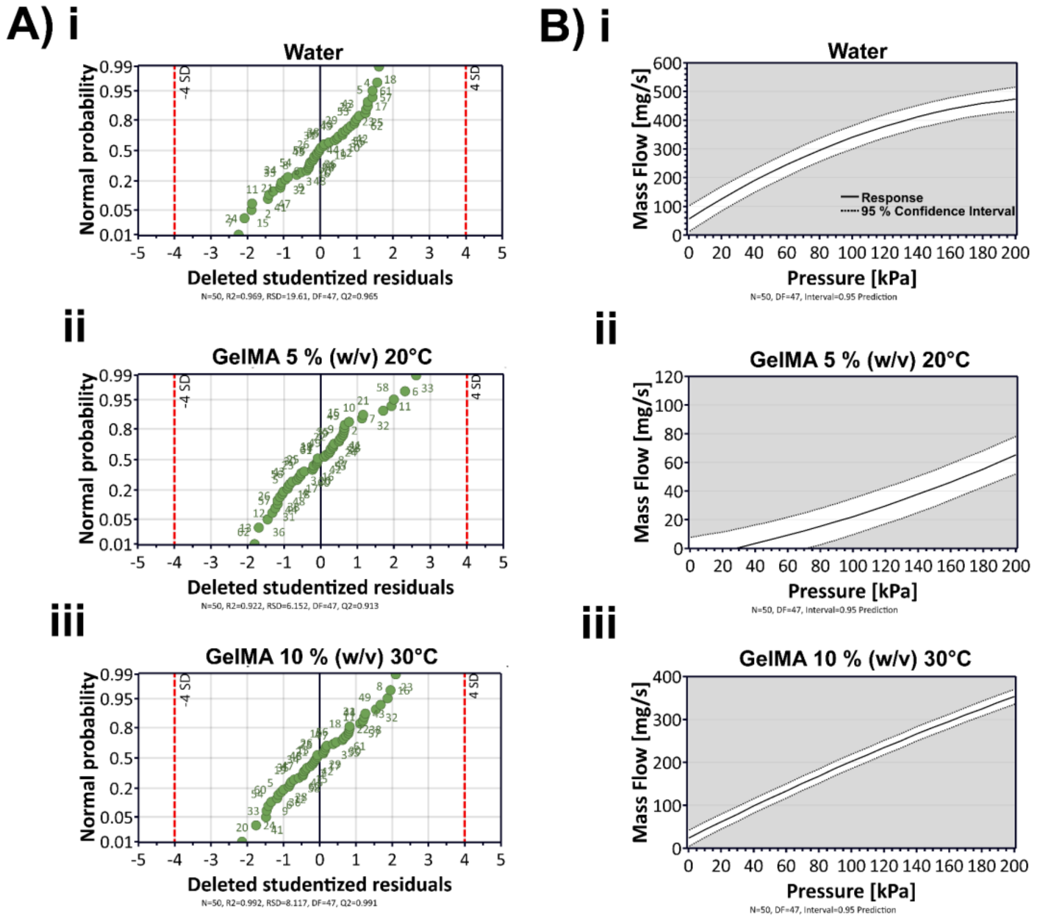


**Figure S3: MODDE® Design of Experiments (DoE) mass flow prediction model.** A) Residuals normal probability plots of collected mass flow rates at extrusion pressures ranging from 20 to 200 kPa i) Water ii) GelMA 5 % (w/v) at 20 °C iii) GelMA 10 % (w/v) at 30 °C. B) Prediction plot of mass flow rates at extrusion pressures between 20 to 200 kPa i) Water ii) GelMA 5 % (w/v) at 20 °C iii) GelMA 10 % (w/v) at 30 °C.

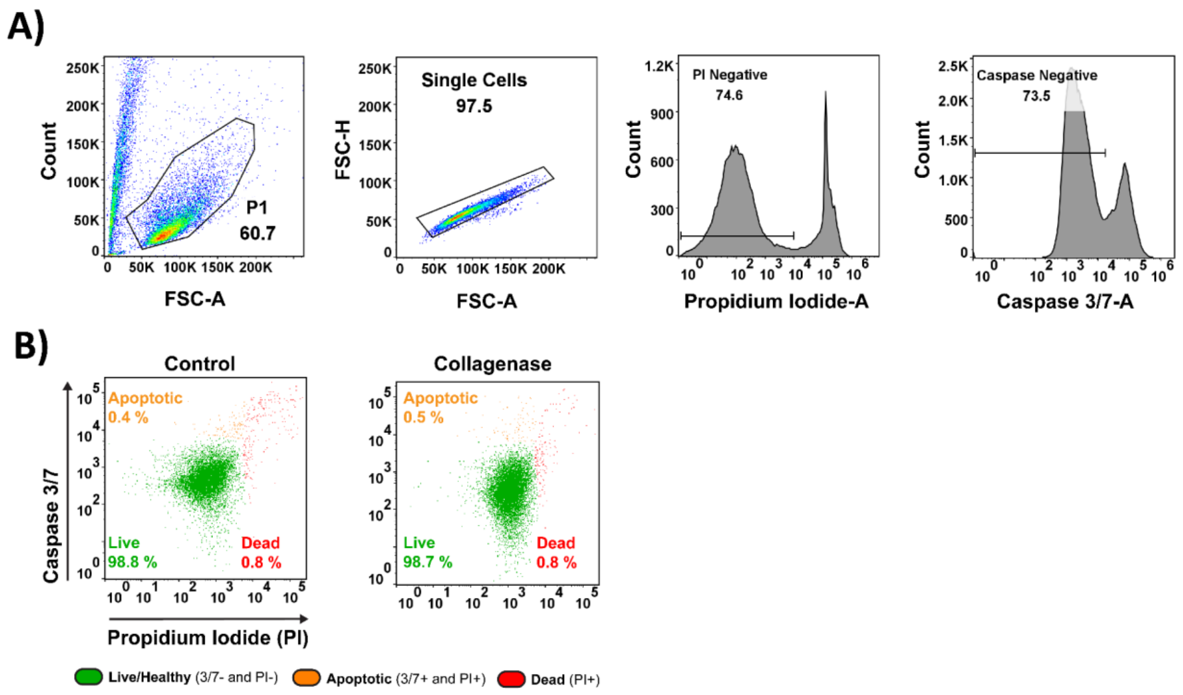


**Figure S1. Flow cytometry gating strategy and the impact of collagenase treatment on cell viability.** A) Gating strategy for immediate shear stress effect assessment, including fluorescence minus one (FMO) controls for Propidium Iodide (PI) and Caspase 3/7 B) Gating strategy for evaluating shear stress effects on crosslinked 5 % (w/v) GelMA after 1 day of culture, with corresponding FMO controls C) Gating strategy for comparing collagenase lysis treatment to control conditions D) Comparative flow cytometry analysis of cell populations following collagenase treatment versus control conditions.

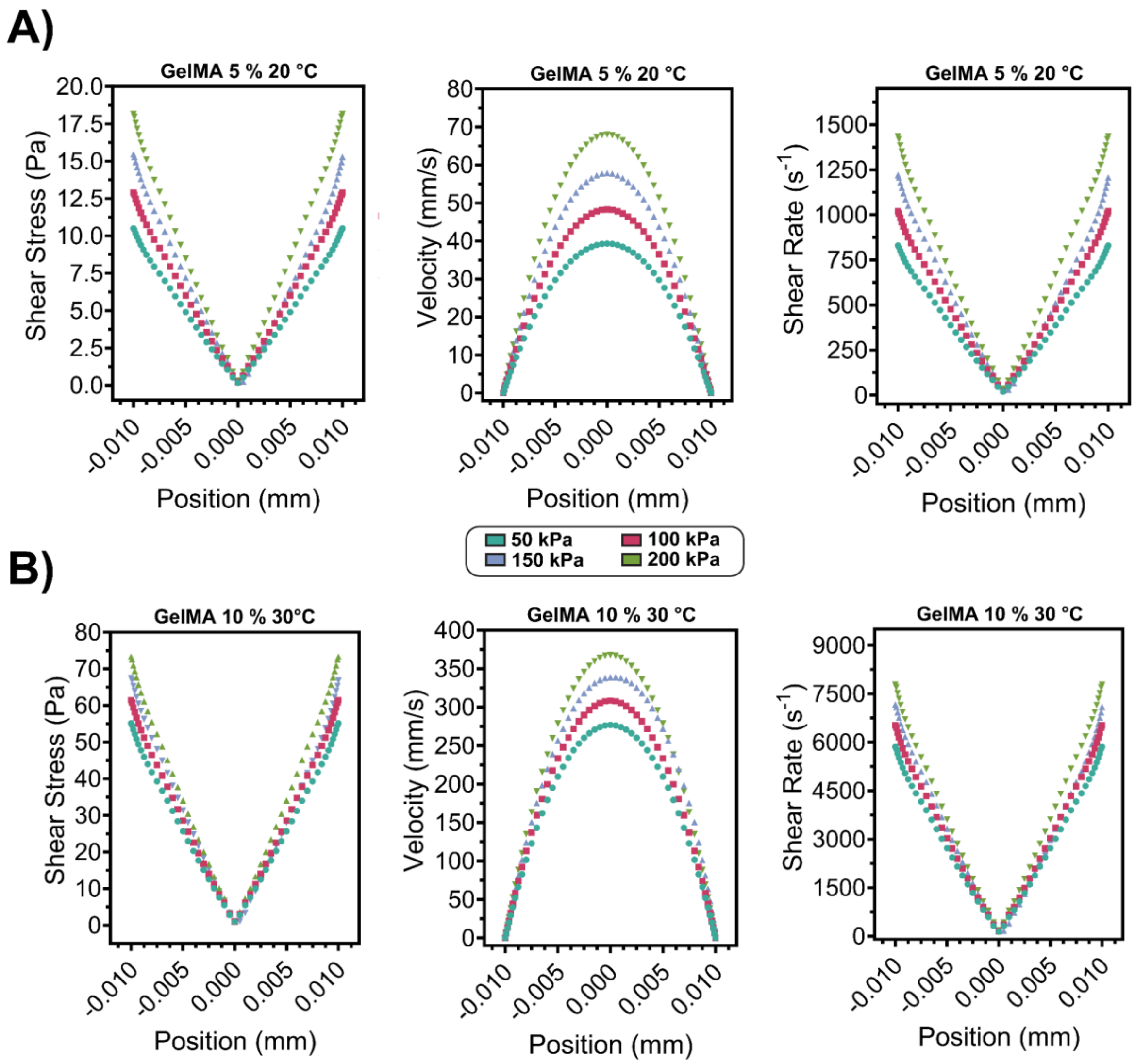


**Figure S5: Simulated 27G nozzle outlet profiles for GelMA 5-10 % (w/v) using Carreau model.** Simulated shear stress (Pa), velocity (mm/s) and shear rate ($s^{-1}$) outlet profiles of A) GelMA 5 % (w/v) at 20 °C and B) GelMA 10 % (w/v) at 30 °C using an extrusion pressure range from 50 to 200 kPa. All simulations utilize an axisymmetric model, contour profiles are mirrored for accurate representation.

**A)**

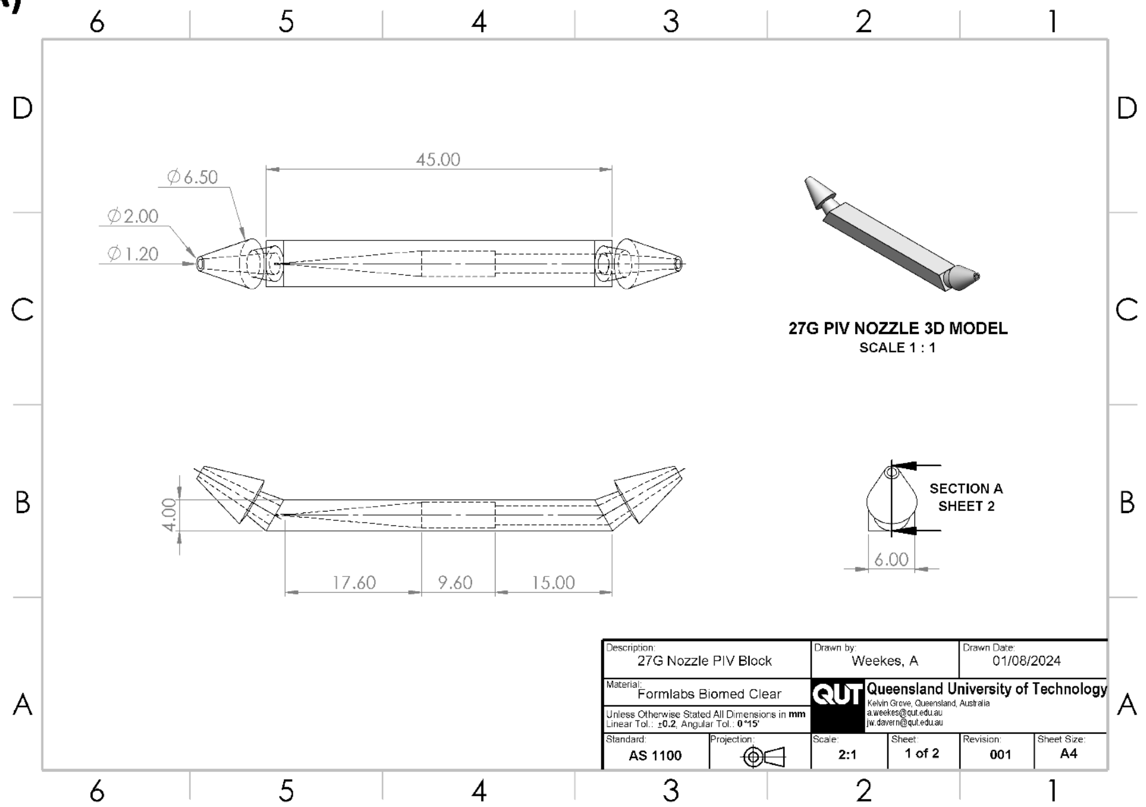


**B)**

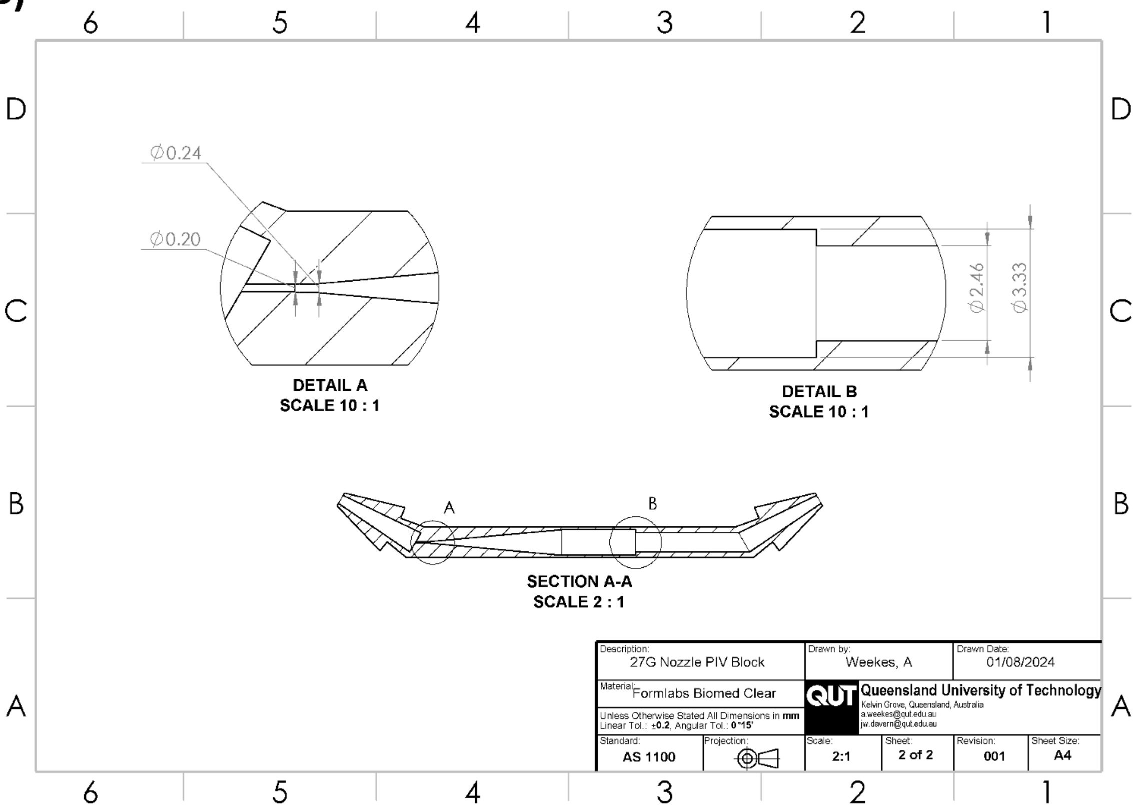


**Figure S6: Engineering drawings of phantom 3D model manufactured for micro-particle velocimetry.** A) Overview of 3D model B) Cross-sectional view of 3D model.
